# KSHV infection of endothelial precursor cells with lymphatic characteristics as a novel model for translational Kaposi’s sarcoma studies

**DOI:** 10.1101/2022.07.25.501362

**Authors:** Krista Tuohinto, Terri A. DiMaio, Elina A. Kiss, Pirjo Laakkonen, Pipsa Saharinen, Tara Karnezis, Michael Lagunoff, Päivi M. Ojala

**Affiliations:** Translational Cancer Medicine Research Program, Faculty of Medicine, University of Helsinki. Finland; Department of Microbiology, University of Washington. Seattle WA, United States of America; Wihuri Research Institute, Biomedicum Helsinki. Finland; Gertrude Biomedical Pty Ltd., Melbourne, Queensland, Australia

**Keywords:** KSHV, HHV-8, lymphatic endothelium, ECFCs, Endothelial precursor cells, Kaposi’s sarcoma, SOX18

## Abstract

Kaposi’s sarcoma herpesvirus (KSHV) is the etiologic agent of Kaposi’s sarcoma (KS), a hyperplasia consisting of enlarged malformed vasculature and spindle-shaped cells, the main proliferative component of KS. While spindle cells express markers of lymphatic and blood endothelium, the origin of spindle cells is unknown. Endothelial precursor cells have been proposed as the source of spindle cells. We previously identified two types of circulating endothelial colony forming cells (ECFCs), ones that expressed markers of blood endothelium and ones that expressed markers of lymphatic endothelium. Here we examined both blood and lymphatic ECFCs infected with KSHV. Lymphatic ECFCs are significantly more susceptible to KSHV infection than the blood ECFCs and maintain the viral episomes during passage in culture while the blood ECFCs lose the viral episome. Only the KSHV-infected lymphatic ECFCs grew to small multicellular colonies in soft agar whereas the infected blood ECFCs and all uninfected ECFCs failed to proliferate. The lymphatic ECFCs express high levels of SOX18, which supported the maintenance of high copy number of KSHV genomes. When implanted subcutaneously into NSG mice, the KSHV-infected lymphatic ECFCs persisted in vivo and recapitulated the phenotype of KS tumor cells with high number of viral genome copies and spindling morphology. These spindle cell hallmarks were significantly reduced when mice were treated with SOX18 inhibitor, SM4. These data suggest that KSHV-infected lymphatic ECFCs can be utilized as a KSHV infection model for in vivo translational studies to test novel inhibitors representing potential treatment modalities for KS.

**Author summary:** Kaposi’s sarcoma herpesvirus (KSHV) is the etiologic agent of Kaposi’s sarcoma (KS). The main proliferative component of KS, spindle cells, express markers of lymphatic and blood endothelium. Endothelial precursor cells, which are circulating endothelial colony forming cells (ECFCs), have been proposed as the source of spindle cells. Here we examined both blood and lymphatic ECFCs infected with KSHV. Lymphatic ECFCs are readily infected by KSHV, maintain the viral episomes and show minimal transformation of the cells, which the infected blood ECFCs and all uninfected ECFCs failed to show. The lymphatic ECFCs express SOX18, which supported the maintenance of high copy numbers of KSHV genomes. The KSHV-infected lymphatic ECFCs persisted in vivo and recapitulated the phenotype of KS tumor cells such as high number of viral genome copies and spindling morphology. These KS tumor cell hallmarks were significantly reduced by SOX18 chemical inhibition using a small molecule SM4 treatment. These data suggest that KSHV-infected lymphatic ECFCs could be the progenitors of KS spindle cells and are a promising model for the translational studies to develop new therapies for KS.

## Introduction

Kaposi’s sarcoma herpesvirus (KSHV) is a gamma herpesvirus and the etiologic agent of Kaposi’s sarcoma (KS) as well as rare B-cell proliferative diseases primary effusion lymphoma and AIDS-associated Castleman’s disease. KS is a hyperplasia consisting of enlarged malformed vasculature and spindle-shaped cells which are the main proliferative component of KS. Spindle cells express markers consistent with an endothelial cell origin [1–5].

Endothelial cells are largely divided into blood vascular and lymphatic endothelial cells. While closely related, the cells making up the blood and lymphatic vasculatures have distinct functions and express specific markers. KS spindle cells express several markers specific to lymphatic endothelium though they also express markers of blood endothelium. However, it has been shown that KSHV infection of endothelial cells in culture induce changes in endothelial cell marker expression. Specifically, KSHV infection of blood vascular endothelial cells (BEC) induces changes consistent with lymphatic differentiation including induction of PROX1 and vascular endothelial growth factor 3, VEGFR3 expression [6–8]. However, infection of lymphatic endothelial cells (LEC) induces the expression of VEGFR1, consistent with its expression in BEC. Therefore, the contribution of these cell types to development of KS tumors is unclear. Another feature of spindle cells is the expression of mesenchymal cell markers, an observation that has prompted the idea that undifferentiated mesenchymal cells might be a cell type of origin [9]. This hypothesis remains elusive since the mesenchymal phenotypes could also emerge due to Endothelial-to-Mesenchymal-Transition (EndMT) induced by KSHV-infection of LECs [10,11].

KS tumors develop primarily in the skin of patients with classic KS. More aggressive AIDS-associated KS can infiltrate internal organs. However, KS is not found in tissues lacking lymphatic vasculature, such as parenchyma and retina. This suggests that LECs are a necessary component of KS development. Furthermore, we and others have previously found that LECs are more susceptible than BECs to KSHV infection [8,12–15]. The KSHV-infected primary human LECs (K-LECs) display a unique infection program characterized by maintenance of high KSHV genome copies, spontaneous lytic gene expression and release of significant amounts of infectious virus [12,13]. This is in striking contrast to BECs and other KSHV-infected in vitro cell models. KSHV infection of neonatal LECs but not BECs allows them to bypass replicative senescence [14]. However, KSHV-infected LECs, like KS spindle cells, are not fully immortalized. Whether LECs are a reservoir for KSHV *in vivo* has yet to be determined.

Endothelial colony forming cells (ECFCs) are circulating cells thought to be endothelial precursors. Previous studies have shown that circulating endothelial precursor cells home to sites of neoangiogenesis [16,17]. However, the contribution of precursor cells to neoangiogenesis is unclear. Previous investigations suggest that circulating endothelial precursors could be important for the development of KS lesions. One study found that KS lesion formation following kidney transplant occurred in distal extremities [18]. Importantly, the spindle cells of the KS tumors were gender matched to the transplanted tissue, not the recipient, suggesting the presence of circulating cells harboring KSHV. Further studies have isolated ECFCs from patients with classic KS, which were found to be positive for KSHV DNA [19]. Interestingly, KSHV infection of endothelial progenitor cells isolated from umbilical cord blood reduces their angiogenic potential [20]. It has been previously proposed that KSHV infected ECFCs could be the source of spindle cells seeding the KS tumors [19–21].

We previously isolated ECFCs from whole blood and found that there are two types of ECFCs [22]. The predominant ECFC expressed markers of blood endothelial cells as has been previously described [23– 25]. Importantly, we identified rare ECFC isolates expressing markers of the lymphatic vasculature. Our recent results show that the key developmental lymphatic transcription factor (TF), SOX18, is expressed in KS tumors and needed to support a unique KSHV infection program with high number of intracellular KSHV genome copies in infected human LECs. SOX18 binds to the origins of KSHV replication and increases the intracellular viral DNA genome copies [12]. Depletion by RNAi or specific inhibition of SOX18 homo- and heterodimerization by a small molecule inhibitor SM4 or R-propranolol [26,27] dramatically decreases both intracellular viral genome copies and release of infectious virus, suggesting SOX18 as an attractive therapeutic target for KS.

Here, we show that similar to LECs, lymphatic ECFCs are more susceptible to KSHV infection and maintain the viral episome in contrast to blood ECFCs. KSHV infection of lymphatic ECFCs allowed proliferation, albeit limited, of these cells in soft agar, suggesting that KSHV may promote enhanced survival of lymphatic ECFCs. We further utilized the SOX18-expressing lymphatic ECFCs as a physiologically relevant KSHV infection model for testing the translational potential of SOX18 small molecule inhibitor SM4 in vitro and in vivo. Together, these data suggest that circulating lymphatic ECFCs could potentially represent virus reservoirs and putative precursors as the source for the initiation of KS tumors.

## Results

### Lymphatic ECFCs are more permissive to KSHV infection than blood ECFCs

We have previously shown that neonatal LECs are more susceptible to KSHV infection than BECs [12,14]. Also, we previously identified two types of ECFCs, ones that expressed markers of BEC and ones that expressed markers of LEC. To determine whether also these lymphatic ECFCs are more susceptible to KSHV-infection than blood ECFCs, we seeded both cell types onto 6-well plates, allowed them to adhere, and followed by infection with identical dilutions of KSHV at either a high or low MOI. After two days, we harvested the cells and performed immunofluorescence for viral latency-associated nuclear antigen (LANA) expression and counted the percentage of cells that were latently infected. As shown in Fig 1A, at high MOI, 98% of lymphatic ECFCs are infected, while only 87% of the blood ECFCs are infected. At high MOI, each cell may have multiple instances of infection, therefore counting the percentage of LANA+ cells may miss cases of superinfection. To address this, we performed the same experiment with a low MOI. Fig 1B shows that lymphatic ECFCs are infected at almost three times the rate of blood ECFCs.

**Figure 1.**
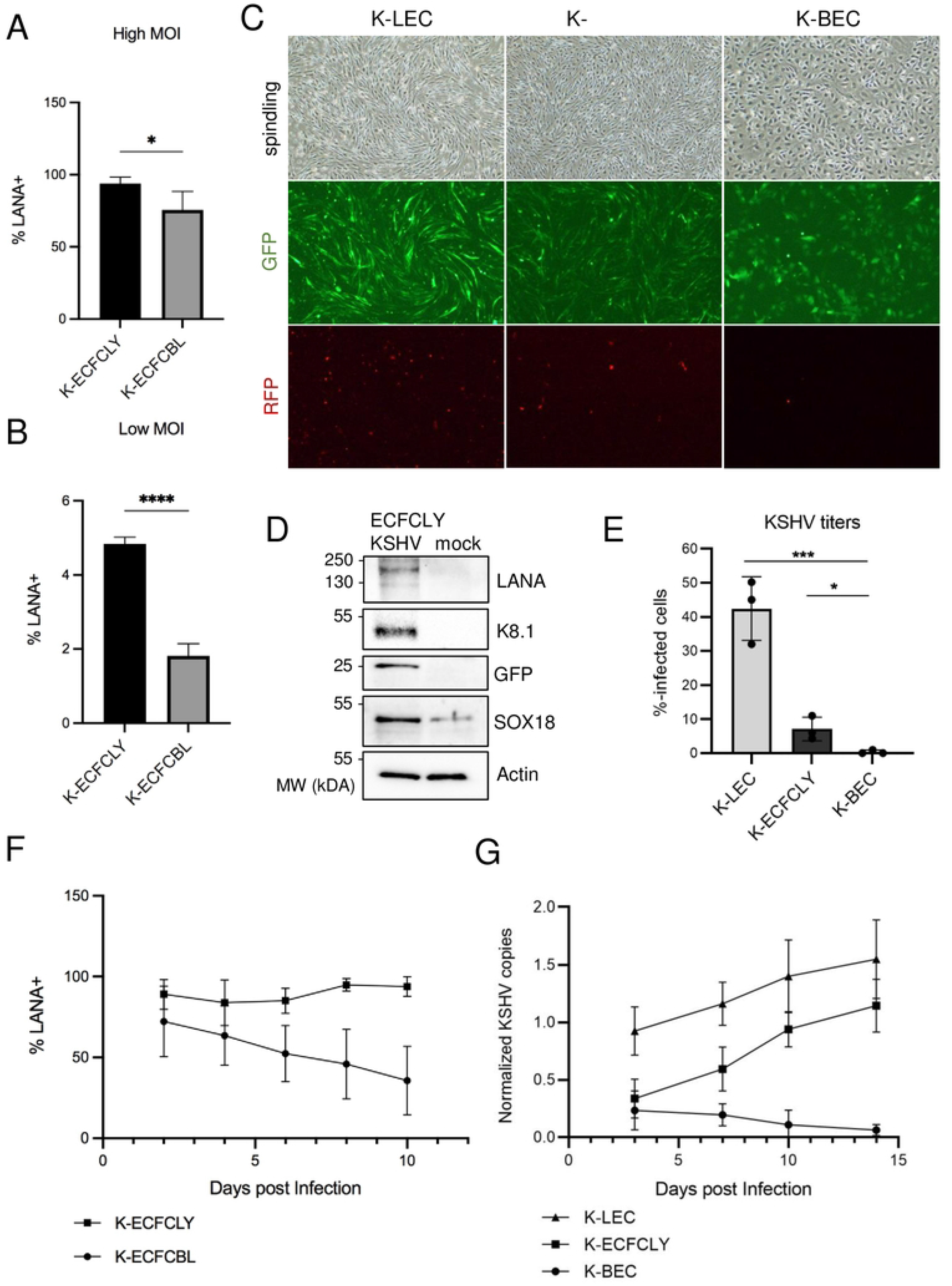
Lymphatic ECFCs are more permissive to KSHV infection than blood ECFCs and maintain the KSHV viral episome. **A**. Lymphatic and blood ECFCs were infected with KSHV (K-ECFCLY and K-ECFCBL respectively) at either a high MOI (upper panel) or **B**. a low MOI (lower panel). At 48 h.p.i cells were harvested and stained with anti-LANA antibody for the number of infected cells and DAPI for the number of total cells. Cells were counted and the percentage of LANA+ cells was determined. These experiments were performed with cells isolated from at least two different donors for each cell type. **C**. KSHV-infection of lymphatic ECFCs, LECs and BECs was compared over 14 days and pictures shown at 7 d.p.i of spindling phenotype, latent infection (GFP) and spontaneous lytic replication (indicated by RFP expression). **D**. Expression of KSHV latent (LANA) and lytic (K8.1) proteins, in addition to GFP and SOX18 were analysed from mock- and infected lymphatic ECFCs. **E**. KSHV titers were measured by a virus release assay on naïve U2OS cells. **F**. Lymphatic (squares) and blood (circles) ECFCs were infected with KSHV and harvested every 2 days and stained for LANA expression. The percentage of LANA+ cells was determined. **G**. Lymphatic ECFCs (squares), K-LECs (triangles) and K-BECs (circles) were KSHV-infected at similar MOI and total DNA was collected at 3, 7, 10 and 14 d.p.i. Relative KSHV genome copies were determined. Experiments were performed three times with similar results. * p < 0.05, *** p < 0.001, **** p < 0.0001.

We also isolated ECFCs using a different protocol where the cells were isolated from the whole blood of four, additional healthy donors, but as an adherent subpopulations rather than from single cell colonies. By fluorescence associated cell sorting (FACS) analyses these cells expressed substantial levels of CD34, VEGFR3, podoplanin, CD31/PECAM1, PROX1 and SOX18, suggesting that they also represent predominantly lymphatic ECFCs (S1 Fig), not blood vascular endothelial ECFCs. Next, we analyzed if these lymphatic ECFCs could support spontaneous lytic replication and production of new infectious viruses, unique to KSHV-infected LECs, but not seen in BECs [12,13]. To this end, we compared the infection of primary human LECs, BECs and the lymphatic ECFCs with recombinant rKSHV.219 [28] on 6-well plates and the infection was followed up for 2 weeks. Fig 1C shows latent infection (indicated by GFP expression) in all cell types, however, the spindling phenotype became visible in 7 days post infection only in lymphatic ECFCs (K-ECFCLY) and K-LECs, but not in K-BECs. Spontaneous lytic replication (indicated by RFP expression) was observed both in K-ECFCLYs and K-LECs, albeit more prominent in K-LECs but not seen in K-BECs. Accordingly, as shown in Fig 1D, in addition to GFP and the latent protein LANA, K-ECFCLY expressed the viral lytic protein K8.1. We have previously shown that K8.1 is prominently expressed by K-LECs, but not in K-BECs [12]. Accordingly, K-BECs did not produce infectious virus, whereas K-ECFCLYs did albeit significantly less than K-LECs (Fig 1E). Similar results showing that K-ECFCLYs can produce infectious virus were obtained when using lymphatic ECFCs from three additional donors (S2 Fig). This indicates that lymphatic ECFCs are permissive to KSHV infection and capable to support productive, lytic replication.

### Lymphatic ECFCs maintain the KSHV viral episome

The KSHV genome is maintained as an episome in dividing cells, though this process is not robust, and the episome is lost over time in cultured endothelial cells [29–32]. Our previous data with neonatal ECs showed that LECs were able to maintain the KSHV episome but BECs lost the genome relatively rapidly over the course of cell passaging [12,14]. To test whether the rate of loss is different between blood and lymphatic ECFCs, we infected cells with KSHV and harvested them every two days for immunofluorescence for LANA expression. Because lymphatic ECFCs are more susceptible to KSHV infection, we used higher MOI to the blood ECFCs to achieve similar initial infection rates. By using LANA staining as a surrogate marker for genome copies, Fig 1F shows that, although in the beginning very similar infection rates were observed for both blood and lymphatic ECFCs, the blood ECFCs gradually lose the episome and by ten days post infection fewer than 40% of cells exhibited punctate LANA staining. In contrast, the lymphatic ECFCs maintained infection at almost 100%.

Additionally, primary LECs and BECs and the lymphatic ECFCs isolated as subpopulation were infected with KSHV using low MOI and harvested for total DNA isolation at days 3, 7, 10 and 14 post infection. Quantification of KSHV genome copies demonstrates that lymphatic ECFCs, similar to LECs, maintain the viral episomes (Fig 1G). Interestingly, K-ECFCLYs and K-LECs show even an increase in genome copies until day 14. In contrast, by using low MOI, K-BECs achieve a lower number of genome copies which are gradually lost within two weeks after infection (Fig 1G).

### KSHV inhibits proliferation but not tube formation of ECFCs

To determine if KSHV confers a growth advantage to ECFCs, we mock-or KSHV-infected blood and lymphatic ECFCs and seeded 3×10^4^ cells 48 hours post infection in 6-well plates and monitored their confluence over time using the live-cell Essen Bioscience IncuCyte imaging system. Fig 2A shows that mock-infected blood ECFCs proliferated at a slightly higher rate than mock-infected lymphatic ECFCs. Interestingly, KSHV infection significantly reduced proliferation of both blood and lymphatic ECFCs. However, there are no significant differences in the reduction of proliferation between these two cell types suggesting that KSHV has a mild inhibitory effect on cell proliferation, but this is not dependent on blood versus lymphatic cell origin. We then also compared the proliferation rate of KSHV-infected and non-infected lymphatic ECFCs that had been isolated as a subpopulation by seeding cells to 96-well plates and subjected them to an EdU pulse for 4h before fixation. The Click-IT EdU staining, measured only from LANA+ cells, indicated only slight difference between the percentage of uninfected and the KSHV-infected lymphatic ECFCs undergoing proliferation (Fig 2B), confirming the mild inhibitory effect on cell proliferation.

**Figure 2.**
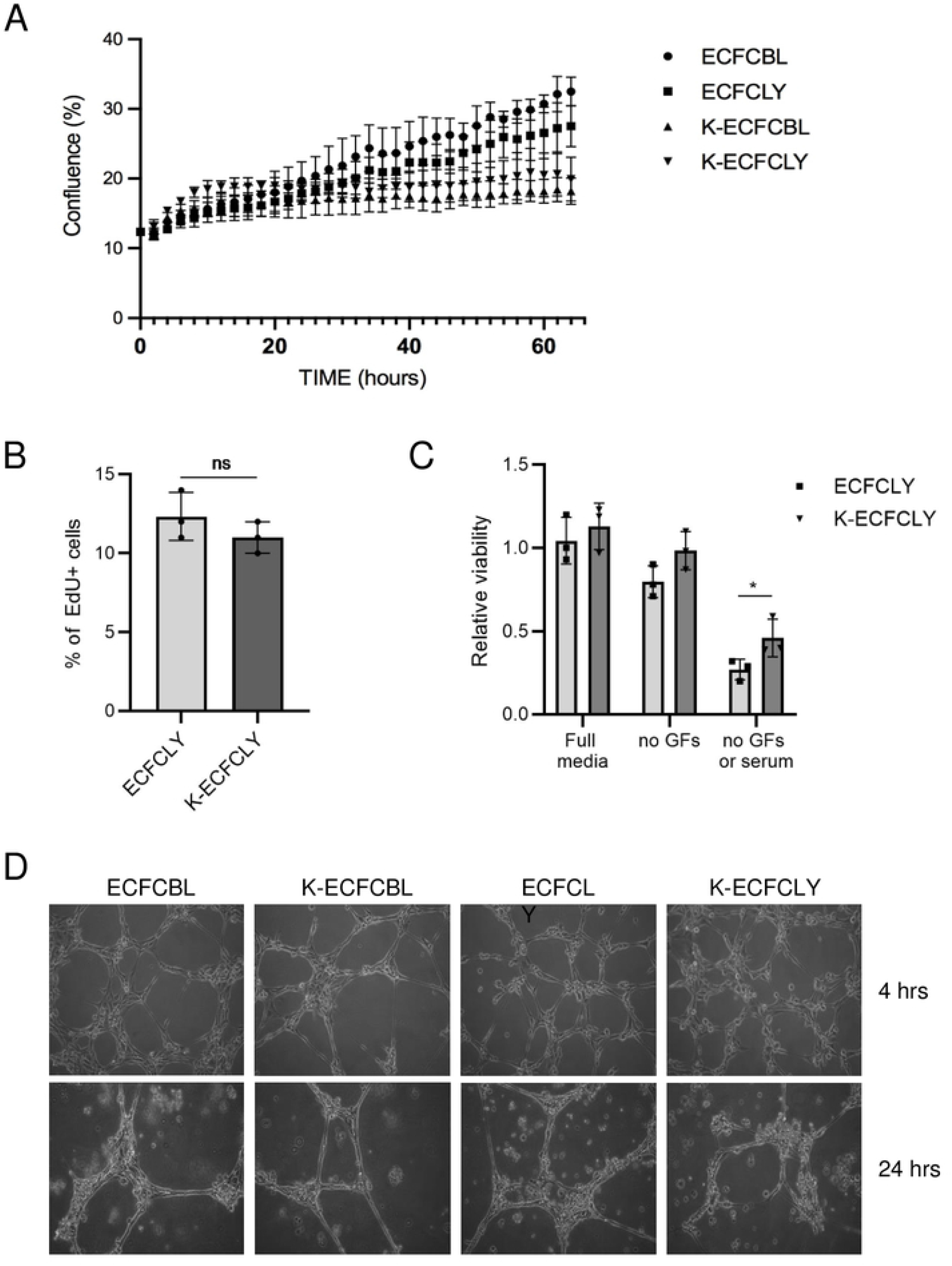
KSHV inhibits proliferation but not capillary morphogenesis. **A**. 48 hours after infection with KSHV, ECFCLY (squares), ECFCBL (circles), K-ECFCLY (upside down triangles) and K-ECFCBL (triangles), were seeded in 6-well dishes and placed in an Essen Biosciences IncuCyte. Pictures were taken every 2 hours and percentage confluence was determined. **B, C**. Mock- or KSHV-infected lymphatic ECFCs were seeded 5 d.p.i on 96-well dishes and **B**. treated with EdU for 4h before fixing. EdU-positive cells were stained, and percentages of proliferating cells were determined. **C**. Relative viability after 7 days in full or serum and/or growth factor deprived media was measured with CellTiter-Glo luminescence assay. **D**. Blood and lymphatic ECFCs were mock- or KSHV-infected. 48 h.p.i cells were plated on Matrigel and monitored for capillary morphogenesis at 4 and 24 hours post plating. * p < 0.05.

To determine if KSHV-infection supports viability of lymphatic ECFCs under limited growth factor conditions, mock and KSHV-infected cells were seeded on 96-well plates. The cells were grown either in normal media containing all the supplements, growth factor reduced media or media without any supplements or serum for 7-day period (media replenished at day 4). As shown in Fig 2C, KSHV-infected lymphatic ECFCs survived better than the mock cells under both the growth factor reduced and serum deprived conditions, indicating increased viability in limiting growth media.

KS tumors are highly vascularized with irregular, enlarged vascular slits, indicating increased neovascularization. We have previously shown that KSHV infection of ECs promotes the stability of capillary networks in a Matrigel model of endothelial cell migration [33]. To measure whether KSHV alters the ability of blood or lymphatic ECFCs to undergo angiogenesis, we mock- or KSHV-infected blood and lymphatic ECFCs. Cells were plated on a Matrigel matrix 48 hours post infection. Fig 2D shows that both blood and lymphatic ECFCs organized into networks on Matrigel at 4 hours post plating (top panels). Interestingly, unlike mature endothelial cells, both blood and lymphatic ECFCs maintained the cord structures at 24 hours post plating (bottom panels). Under these conditions, KSHV had little effect on the ability of ECFCs to organize on Matrigel and maintain their capillary stability.

### KSHV confers a survival advantage to lymphatic ECFCs grown in soft agar

We next examined if KSHV promotes anchorage independent growth of ECFCs in soft agar, an indicator of cell transformation. We mock- or KSHV-infected blood and lymphatic ECFCs and, after 48 hours, embedded a single cell suspension in a soft agar matrix which was overlaid by normal growth media. We then monitored the growth and formation of small colonies of cells over the course of four weeks. Fig 3A shows images of cell suspensions following four to five weeks of growth. We could not find any evidence of multicellular colonies in the uninfected blood nor lymphatic ECFCs. Only single cells could be visualized even when incubated 36 days indicating these cells are unable to proliferate in the absence of attachment to a solid surface. Additionally, KSHV-infected blood ECFCs (K-ECFCBL) showed no indication of multicellular colonies in soft agar. Interestingly, after four weeks in soft agar culture, many multi-cellular colonies were found in the wells containing K-ECFCLYs indicating the ability of KSHV infected lymphatic ECFCs to grow in an anchorage independent fashion. Scanning and quantifying multiple experiments indicated that approximately 10% of the KSHV infected ECFCLYs formed multicellular colonies (Fig 3B).

**Figure 3.**
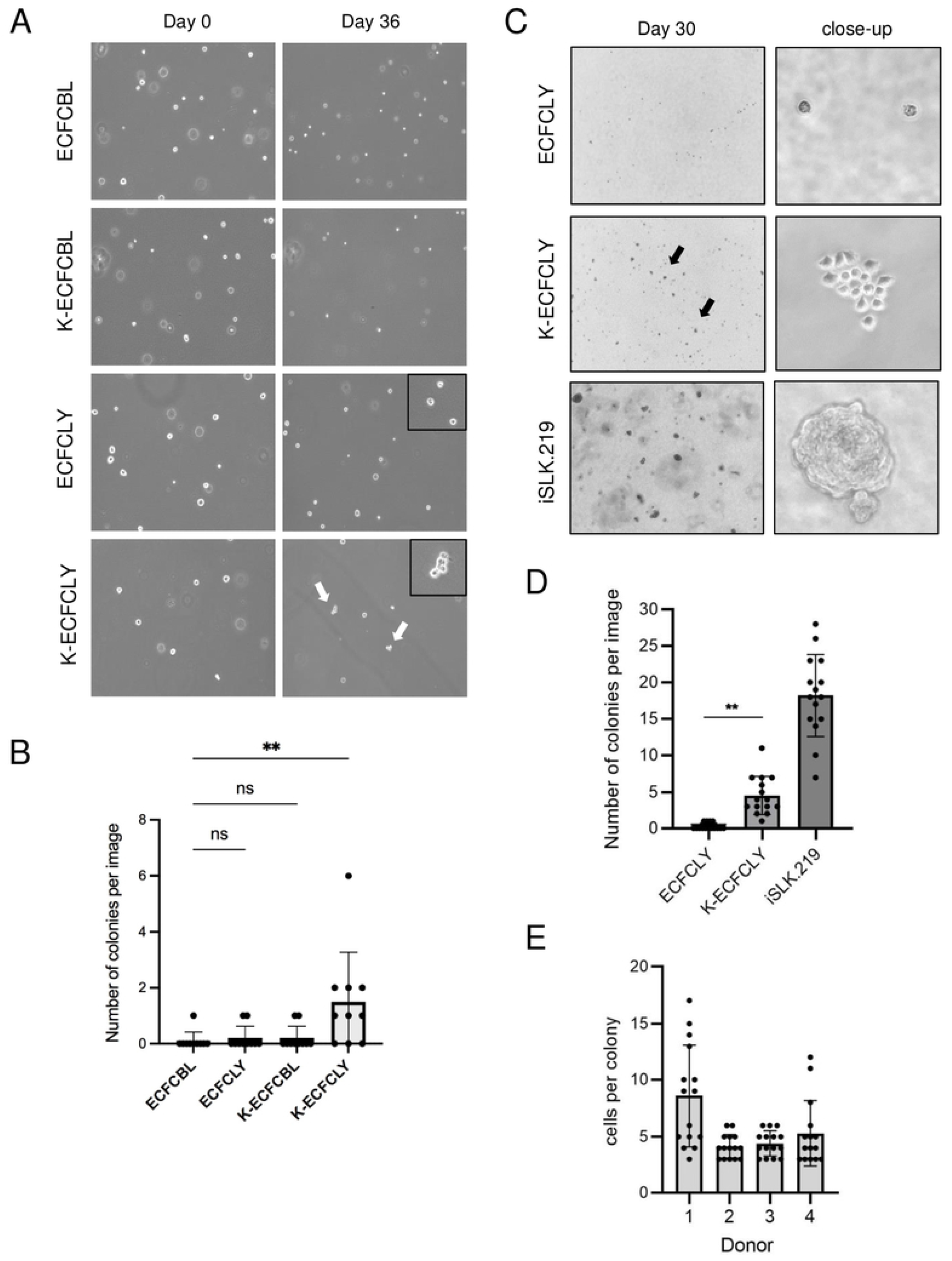
KSHV confers a survival advantage to lymphatic ECFCs grown in soft agar. **A**. Mock or KSHV-infected lymphatic and blood ECFCs were embedded 48 h.p.i in soft agar overlaid with 0.5ml growth media, and media was replenished every 2-4 days. **B**. The number of colonies with two or more cells at 36 days post plating were quantified. These experiments were performed up to eight times with similar results. **C**. Lymphatic ECFCs mock- or KSHV-infected were embedded 5 d.p.i. in soft agar. iSLK.219 was used as a positive control cell line. Quantification of number of colonies of at least 3 cells **(D)** and number of cells in colonies from 4 different donors **(E)**. Experiments were performed at least three times with similar results. ** p < 0.01.

The capacity of KSHV-infected cells to grow in soft agar was also addressed using the lymphatic ECFCs isolated as an adherent subpopulation. To this end, non-infected and KSHV-infected lymphatic ECFCs from four different donors were embedded as single cells to soft agar seven days post infection. Fully transformed renal cell carcinoma cell line stably infected with KSHV, iSLK.219 [34], was used as a positive control. We then monitored the growth and formation of small colonies of cells over the course of four weeks. Fig 3C shows images of cells from donor 1 after four weeks of growth. The colonies were calculated and quantified as shown in Fig 3D-E. We could not detect any evidence of multicellular colonies in the non-infected samples, however, after four weeks in soft agar, many colonies were observed in the wells containing K-ECFCLYs. These results further support the ability of KSHV infected lymphatic ECFCs to proliferate in an anchorage independent fashion.

Together, this suggests that KSHV may promote the survival and proliferation of lymphatic ECFCs but not blood ECFCs in soft agar. However, this effect was limited to small colonies containing approximately 3-17 cells. Fully transformed cells like iSLK.219 form robust colonies with 100s of cells (Fig 3C-D). Therefore, KSHV allows significant but limited proliferation of cells in soft agar indicating minimal transformation of the cells. The slow expansion of K-ECFCLYs in soft agar as compared to the iSLK.219 cell line would be consistent with the slower growth of KS tumors compared to many more aggressive tumors like renal cell carcinoma.

### Gene expression changes induced by KSHV infection of lymphatic ECFCs

To determine the effects of KSHV infection on blood and lymphatic ECFCs, we measured the global gene expression differences between these cells upon primary KSHV infection. mRNA from blood and lymphatic ECFCs that were mock- or KSHV-infected, was analyzed by high throughput RNA sequencing 48 hours post infection. We also compared these gene expression profiles to previous data we generated from mature BECs and LECs [14]. S3A Fig shows a Venn diagram comparing all genes upregulated during KSHV infection by at least 2-fold in all cell types. Approximately 125 genes are specific to KSHV-infected BECs (both mature and ECFCs), while 52 genes were specific to LECs. To validate the RNA-sequencing results, we analyzed selected genes that had differential expression between blood and lymphatic ECFCs using custom PrimePCR plates (S3B Fig). These data are consistent with the gene expression changes identified by high-throughput sequencing.

Using Cytoscape software and the Gene Ontology classification application BINGO, we determined the Gene Ontology terms that were highly enriched among the K-ECFCBLs and K-ECFCLYs. The most highly enriched categories for blood ECFCs are listed in S1 Table. Interestingly, these include many immune response categories. The individual genes for some of these categories are listed in S2 Table. In contrast, lymphatic ECFCs were not enriched in immune response categories, but were enriched in genes involved in cell differentiation and signaling (S3 & S4 Tables). To confirm whether genes involved in the innate immune response are upregulated in KSHV-infected blood ECFCs but not lymphatic ECFCs, we performed real-time quantitative RT-PCR on RNA from mock and KSHV infected blood and lymphatic ECFCs. S3C Fig shows that several genes involved in innate immunity, such as viperin and IFI6, are induced by KSHV in blood ECFCs but not in lymphatic ECFCs. This suggests that lymphatic and blood ECFCs may have differential innate immune responses to KSHV infection.

High-throughput RNA sequencing was repeated with lymphatic ECFCs, isolated as a subpopulation from a different donor. Cells were mock- or KSHV-infected for seven days before total RNA was harvested for sequencing. The gene expression profiles were compared to the data obtained from lymphatic ECFCs infected with KSHV for 48h as described above. S3D Fig shows a Venn diagram of 1444 shared gene expression changes between the two infected ECFCLY-isolates, also listed in S6 Table. The higher number of differentially expressed genes with ECFCLYs infected for seven days is most likely due to the longer exposure to virus thus allowing more phenotypic changes to occur.

S3E Fig shows a Volcano blot of differentially expressed genes between mock and KSHV-infected ECFCLY after 7 days of infection. Interestingly, significantly upregulated genes include those involved in lymphatic endothelial specification, such as LYVE1, Podoplanin, and VEGFC. Also, some mesenchymal genes implicated in EndMT were moderately, but significantly, upregulated. Selected genes were validated with RT-qPCR shown in S3F Fig, and they are consistent with the gene expression changes identified by high-throughput sequencing.

### SOX18 inhibition reduces the hallmarks of KSHV infection in lymphatic ECFCs in vitro and in vivo

Our recent data demonstrated that SOX18, expressed abundantly in KS tumors and K-LECs, binds to the origins of KSHV replication and increases the intracellular viral DNA genome copies [12]. To address the SOX18 role in K-ECFCLYs the cells were treated with siRNA targeting SOX18 or non-targeting control siRNA (siNeg) after five days of infection and analyzed three days later for KSHV genome copies, number of LANA positive cells and infectious virus release on naïve U2OS cells (Fig 4A-D). Similar to K-LECs, depletion of SOX18 significantly reduced the intracellular viral genome copies, number of LANA-positive cells and the virus titers (Fig 4A-D). To measure the effect of SOX18 inhibition by the SM4 inhibitor on lymphatic K-ECFCLYs, GFP and RFP positive cells (to monitor latent and lytic infection), spindling phenotype of the cells, in addition to the above-mentioned phenotypes were measured as the hallmarks of KSHV infection. K-ECFCLYs (at 5 days p.i.) were treated with SM4 at concentrations 1 µM, 25 µM and 50 µM for 6 days and DMSO was used as a control. Fig 4E shows that similarly to depletion of SOX18 by siRNA, 25 µM of SM4 was sufficient to reduce the GFP positive cells, spindling phenotype and RFP signal indicating cells undergoing lytic cycle. Furthermore, SM4 significantly, and in a dose-dependent manner, reduced the number of KSHV genome copies, LANA positive cells and released infectious virus from the infected lymphatic ECFCs (Fig 4F-H). These data demonstrate that SOX18 is required to maintain the high number of intracellular KSHV genomes and viral titers in the K-ECFCLYs, similar to K-LECs [12].

**Figure 4.**
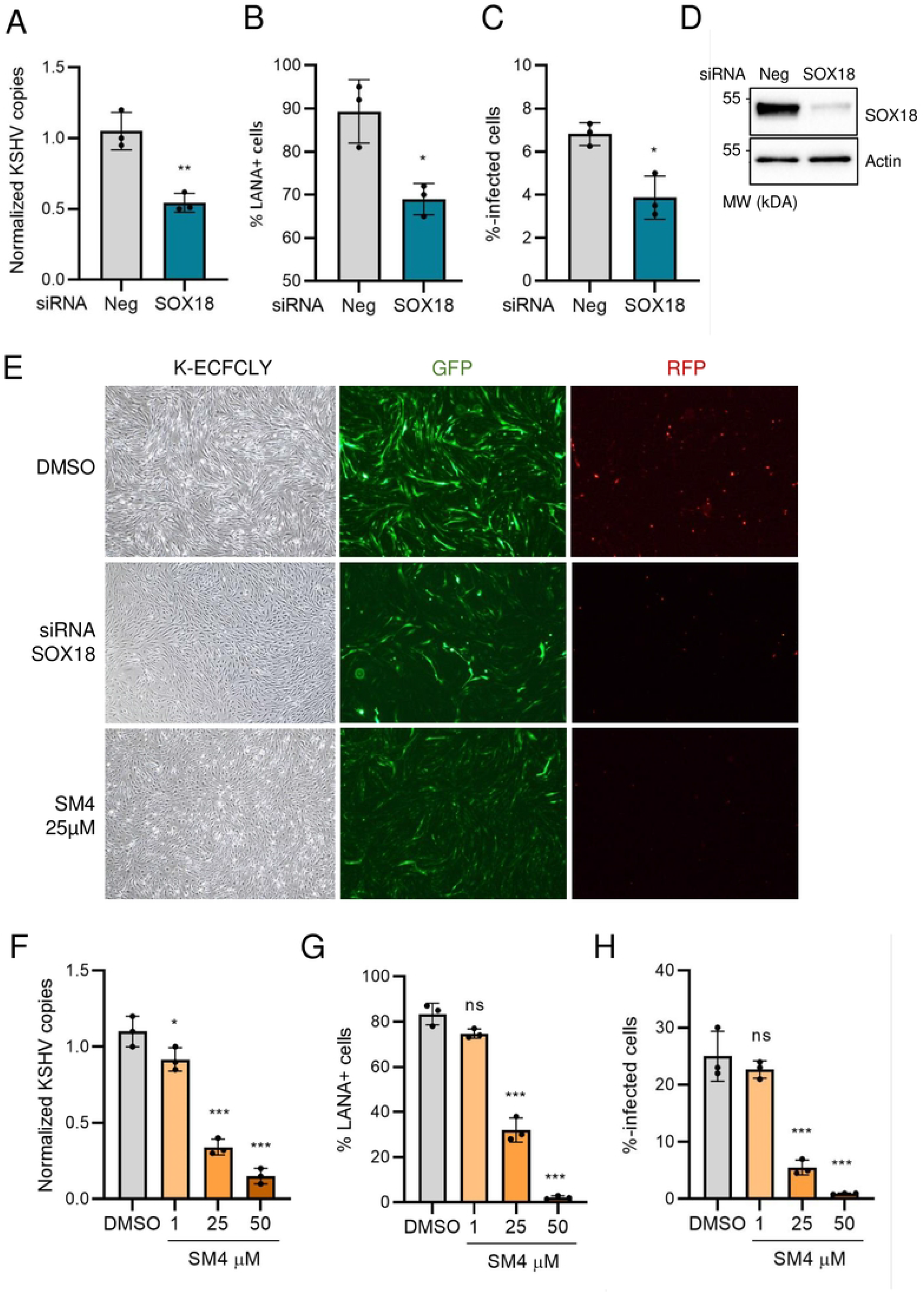
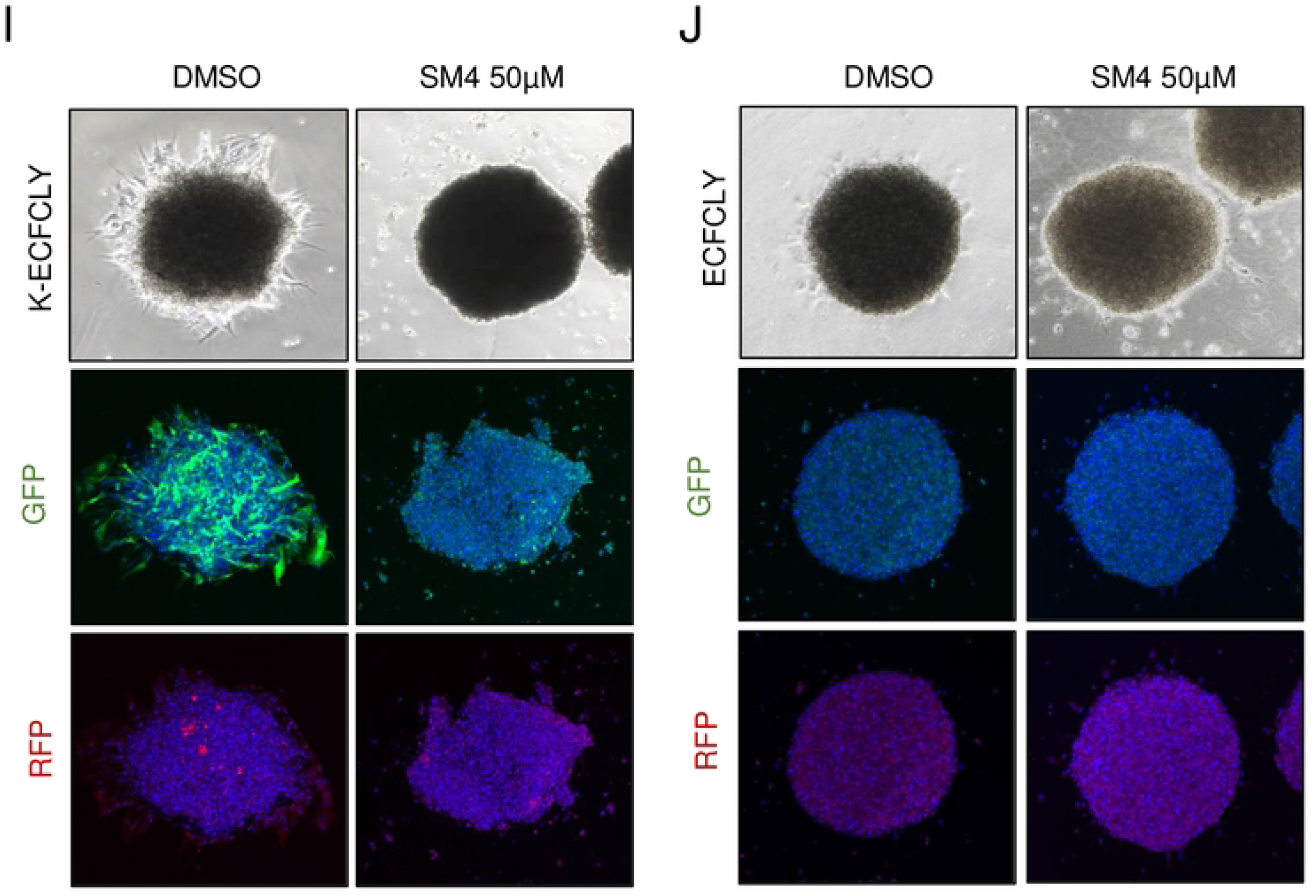
SM4 inhibit the hallmarks of KSHV infection in lymphatic ECFCs in vitro. Lymphatic ECFCs were infected with KSHV for 5 days, and treated with **A-E**. siRNA targeting SOX18, or control siRNA (siNeg) transfected for 72h or **E-H**. with indicated concentrations of SM4 inhibitor or DMSO control for 6 days, replenished at day 3. The effect of treatments is shown as quantified **A, F**. KSHV genome copies from total DNA, **B, G**. percent of LANA positive cells on 96-well plates and **C, H**. KSHV titers measured from virus release assay on naïve U2OS cells. **D**. The efficiency of SOX18 silencing shown at a protein level. **E**. Changes in spindling morphology, GFP (latent) and RFP (lytic) cells shown with siRNA targeting SOX18 or 25 µM of SM4 and DMSO control. **I-J**. Lymphatic ECFCs were either mock or KSHV-infected, allowed to form spheroids overnight and then embedded in 3D fibrin. Spheroids were treated for 6-days with 50 µM of SM4 or DMSO control, and fixed. Phase contrast and confocal images are shown at treatment day 6. * p < 0.05, ** p < 0.01, *** p < 0.001.

By using a three-dimensional (3D), organotypic cell model for KSHV infection we have previously shown that KSHV promotes the ability of mature lymphatic endothelial cells to invade into 3D cross-linked fibrin [10]. Next, we tested if KSHV infection would induce the 3D sprouting growth of the lymphatic ECFCs and if it was dependent on SOX18 function. To this end, mock- and KSHV-infected lymphatic ECFCs cells were first allowed to form spheroids overnight and then embedded in 3D fibrin. As shown in Fig 4I KSHV induced outgrowth of sprouting cells from the lymphatic ECFC spheroids, which was completely abolished during the 6-day 50 µM SM4 treatment. The mock infected lymphatic ECFC spheroids did not sprout and SM4 did not have any effect on their morphology (Fig 4J). Together these data support that the lymphatic ECFC represent a viable model for KSHV infection and testing of the translational potential of novel targets for KS.

### SOX18 inhibition reduces the hallmarks of KSHV infection in vivo

As the KSHV-infected lymphatic ECFCs support the complete lytic replication program, proliferate, and show emerging transformation, we decided to test their long-term persistence *in vivo* in NSG mice when implanted subcutaneously. Mock and KSHV-infected lymphatic ECFCs were implanted subcutaneously in Matrigel after seven days of infection (7 d.p.i.). After a 30-day period, visible cell plugs were collected and analyzed for the presence of KSHV DNA and intensity of GFP (from the rKSHV.219 latent infection) and LANA by IFA from the paraffin embedded sections (Fig 5A). Fig 5B shows that the K-ECFCLYs recapitulate the spindling phenotype of KS tumor cells in the lesions and express LANA suggesting that they represent a promising *in vivo* model to investigate the potential treatment modalities for KS.

**Figure 5.**
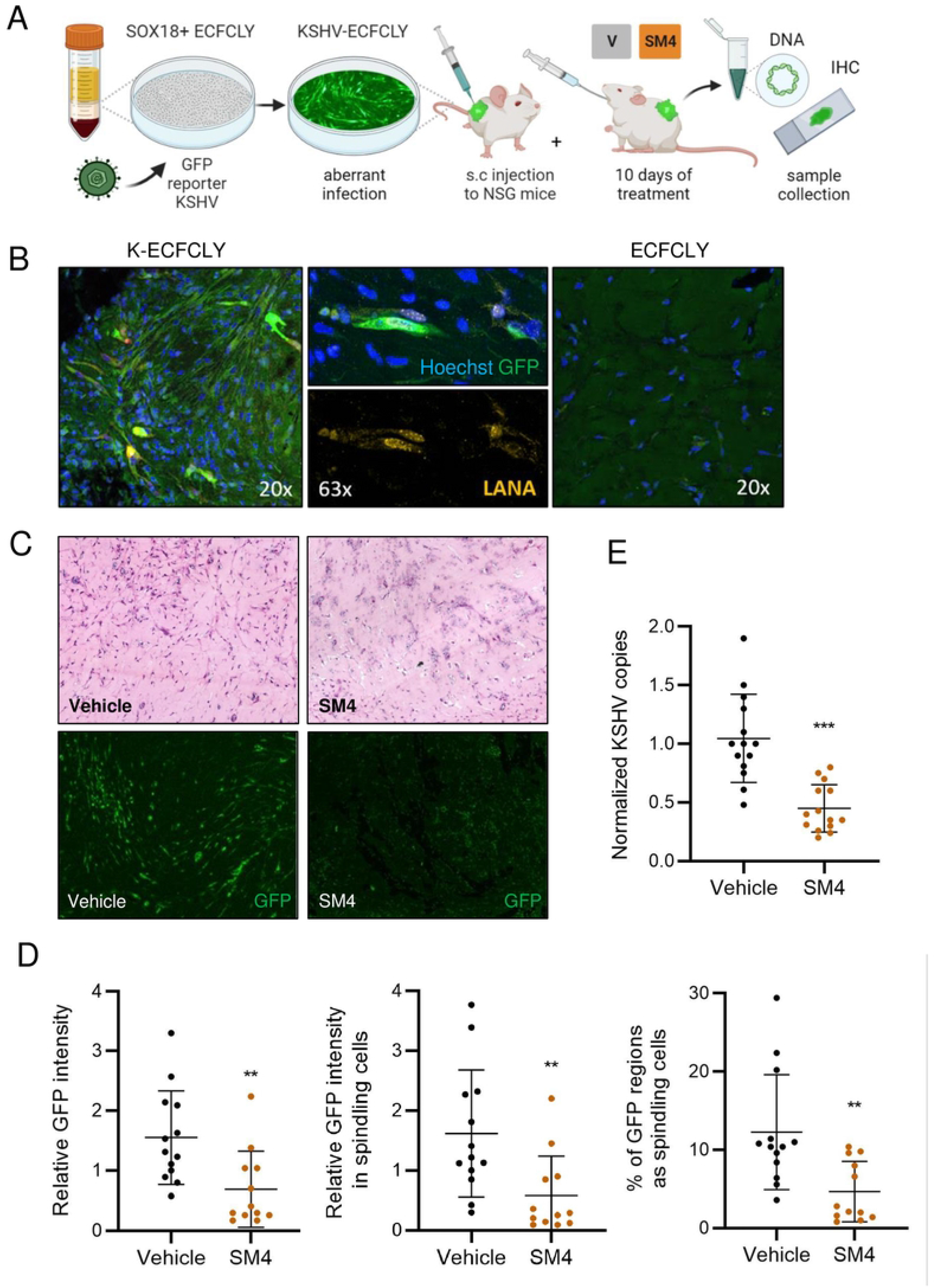
SM4 inhibit the hallmarks of KSHV infection in lymphatic ECFCs in vivo. **A**. Schematic of the experimental in vivo layout of SOX18 inhibition on ECFCLY KSHV-infection model and sample collection. **B**. Lymphatic ECFCs infected with KSHV for 7 days, or left uninfected, were implanted subcutaneously to NSG mice. Long-term persistence of K-ECFCLY seen as expression of spindling GFP-positive cells and staining of nuclear LANA dots, not seen with uninfected ECFCLY, from grafts extracted 30 days post implantation. **C-E**. Lymphatic ECFCs and LECs, infected with KSHV for 7 days, or left uninfected, were implanted subcutaneously to NSG mice with ratio of 90-95% of K-ECFCs and 5-10% of K-LECs. 24h post implantation the mice were treated with either SM4 or Vehicle control daily for 10 days after which mice were sacrificed and grafts were collected for total DNA and IHC. **C**. H&E staining imaged with slide scanner and GFP imaged with confocal microscope. Quantification of **D**. overall GFP intensity, GFP intensity only in spindling cells and spindling morphology (n = 12/ SM4 and 13/ Vehicle group) and **E**. normalized KSHV genome copies from total DNA measured with RT-qPCR between SM4 or Vehicle treated mice (n = 14/group). ** p < 0.01, *** p < 0.001.

We next tested the effect of SM4 on similar model (Fig 5A). To this end, 4×10^^6^ mock- or KSHV infected lymphatic ECFCs, mixed with K-LECs, were injected subcutaneously into NGS mice 7 d.p.i. K-LECs were included to provide more spontaneously lytic cells that can contribute to the inflammatory microenvironment and produce more infectious virus. The final proportions of cells were 90-95% of K-ECFCLYs and 5-10% of K-LECs. The next day SM4 or vehicle (80% PEG-400, 10% Solutol HS-15, 10% dH2O) was administered orally at a dose of 25mg/kg of body weight daily for 10 sequential days as described in [27], after which the cell plugs were collected and analyzed by IHC for cell morphology, IFA for KSHV infection, and qPCR for KSHV DNA.

Interestingly, IHC of cell plugs from mice treated with SM4 displayed several necrotic areas in the grafts, whereas the vehicle group showed cells with a spindle-like, elongated morphology, often found in clusters (Fig 5C, left panels). IFA was performed for sections to distinguish the GFP-expressing, KSHV-infected cells with spindle-like morphology. Stained sections imaged with a confocal microscope are shown in Fig 5C (right panels). The intensity of GFP signal and the coverage of spindling cells in the sections differed significantly between the treatment groups (Fig 5D). While GFP signal in spindling cells was observed in the Vehicle group, the overall GFP signal in the SM4 group was weak, having also significantly less cells with spindling morphology. Quantification showed that the Vehicle group had higher GFP intensity and more spindling cells, when normalized to the section area, and when compared to the SM4 treatment group (Fig 5D). The mock infected lymphatic ECFCs did not express GFP and were used to subtract the autofluorescence for quantification. These results indicate strong potential of SOX18 inhibition to reduce the hallmarks of KSHV infection in this novel in vivo KSHV infection model.

We next measured the KSHV genome copy numbers by qPCR from the extracted total DNA of the grafted samples. The human repetitive ALU-sequences as an internal control were used as a highly sensitive and specific quantification for the human cells among the abundant rodent cells. The KSHV genome copies were quantified using primers specific for LANA. Fig 5E shows that SM4 significantly decreased the relative KSHV genome copy numbers compared to the Vehicle control group further supporting the potential of SOX18 inhibition as a viable therapeutic modality for KS.

## Discussion

Studies of KS tumors are limited by the lack of a clear understanding of the source of the main proliferating tumor cell, the spindle cell. While spindle cells express markers of lymphatic endothelium and most closely align to the gene expression pattern of lymphatic endothelial cells, they also express blood vascular endothelial and mesenchymal cell markers. However, in contrast to angiogenic endothelial cells, quiescent endothelial cells in mature vessels generally do not migrate. There is evidence that the cells that seed KS tumors are able to traffic from the kidney as KS of donor origin formed to the lower extremities of the recipient following a kidney transplant from a KSHV-positive donor [18]. Interestingly, endothelial precursor cells traffic throughout the body through the blood and are known to traffic to sites of angiogenesis. Thus, it has been proposed by multiple groups that endothelial precursor cells are a likely source of KS spindle cells [19–21]. Therefore, we sought to isolate the precursor ECFCs and characterize KSHV infection in those cells. When isolating ECFCs from human blood, we found that the standard endothelial colony forming cells could be separated into isolates that expressed blood endothelial markers and isolates that expressed lymphatic specific markers. We also isolated ECFCs from four donors as an adherent subpopulation without clonal expansion of individual colonies. Interestingly, these isolates represented uniform cultures of predominantly lymphatic ECFCs. This might indicate that lymphatic ECFCs have a growth advantage over the blood ECFCs and take over the culture when cells proliferate during colony formation.

Similar to the neonatal blood and lymphatic endothelial cells isolated from foreskins, the lymphatic ECFCs were more permissive to KSHV infection and the viral episomes were maintained over multiple passages at higher levels as compared to the blood ECFCs. KSHV infection did not alter the ability of either cell type to form capillary-like structures in Matrigel indicating that KSHV does not dramatically alter the angiogenic potential of either cell type. However, infection with KSHV slightly decreased the proliferation rate of both blood and lymphatic ECFCs. We have also noted a decrease in proliferation of neonatal and adult endothelial cells [12], so this appears to be a common phenotype of KSHV infection of endothelial cells in general (DiMaio and Lagunoff, unpublished).

To determine if there was any evidence of transformation of the ECFCs despite the reduced proliferation, we examined the proliferation of non-infected and infected blood and lymphatic ECFCs following detachment from a solid matrix. Interestingly, neither non-infected or infected blood ECFCs proliferated in soft agar nor did the uninfected lymphatic ECFCs. However, in a large number of studies using six different isolates of KSHV-infected ECFCLYs, approximately 10% of the cells formed small colonies following incubation of over four weeks in soft agar. While the colonies did not expand beyond the small 3-17 cell colonies, in all the experiments performed we did not detect any colony formation in the non-infected blood or lymphatic ECFCs or the infected blood ECFCs indicating that this proliferation only occurred in the KSHV-infected lymphatic ECFCs and was reproducible. While there were numerous colonies formed, not every cell formed a colony. Thus, the transformation phenotype was limited while a similar assay with iSLK.219, a renal cell carcinoma cell line stably infected with a recombinant KSHV, led to very large colonies in soft agar of hundreds of cells or more. The iSLK.219 cells also formed colonies more rapidly. Of note, in line with our findings, KS tumors are fairly indolent and do not grow rapidly.

To begin to decipher why there are differences in the behavior of lymphatic and blood ECFCs following KSHV infection we performed RNAseq analysis of non-infected and KSHV-infected lymphatic and blood ECFCs. We identified 758 genes that are enriched in KSHV-infected blood ECFCs compared to lymphatic ECFCs (S5 Table). In contrast, 1,395 genes are enriched in lymphatic ECFCs. Interestingly, when we analyzed the lists of genes for enrichment of Gene Ontology categories, we found that many of the genes induced by KSHV in blood ECFCs were involved in the innate immune response. These include MX1 and MX2, which are interferon-induced genes that are viral restriction factors. MX2 in particular has been shown to be effective at inhibiting herpesviruses, including KSHV [35,36]. Other interferon response genes are also enriched in KSHV-infected blood ECFCs such as IFI6 and RSAD2. It is unclear what effect this enhanced interferon response has on blood ECFCs. However, it could play a role in the reduced susceptibility of blood ECFCs to KSHV infection or the increased rate of episome loss. We previously found that in neonatal LECs STING was not activated [37]. Further work is necessary to determine if the same is true for lymphatic ECFCs. In contrast to the blood ECFCs, genes upregulated by KSHV in lymphatic ECFCs were found to be involved in lymph vessel development and intracellular signaling. What role these genes have on KSHV infection of lymphatic ECFCs remains to be elucidated.

We recently reported high SOX18 expression in KS tumors and that was required to maintain the high number of intracellular KSHV genome copies in infected LECs, suggesting SOX18 as an attractive therapeutic target for KS [12]. Similar to K-LECs we found that KSHV-infected lymphatic ECFCs express high levels of SOX18 and inhibition of SOX18 by a specific small molecule inhibitor SM4 significantly reduced the intracellular viral genome copies also in KSHV-infected EFCFLYs. Since K-LECs do not support long-term growth in culture and do not show even emerging signs of transformation they cannot be used in preclinical in vivo studies to test the efficacy of SOX18 inhibitors. Therefore, we investigated whether KSHV-infected lymphatic ECFCs engrafted into NSG mice could serve as a physiologically relevant infection model to test the potential of SOX18 inhibition as a therapeutic modality for KS. The KSHV-infected ECFCLYs survived as xenografts in mice for at least a month and recapitulated the hallmarks of KSHV infection, which were significantly reduced upon SOX18 inhibition by SM4.

Together our data suggest that lymphatic ECFCs are more susceptible to KSHV infection and may have increased survival and proliferation capabilities during infection. While this does not definitively solve the issue of the source of spindle cells, it shows that infected lymphatic ECFCs are certainly a possible source. Moreover, our data supports that lymphatic ECFCs represent a viable new in vivo model for KSHV infection, suitable for translational studies testing new therapeutic approaches for KS. Further work to analyze KSHV infection of lymphatic ECFCs is warranted.

## Materials and Methods

### Cells

For isolation of ECFCs the blood samples were obtained from the Puget Sound Blood Bank (Seattle, WA, USA) as either outdated whole blood (approximately 500 ml per unit) or leukocyte reduction filters, which yielded between 4-7×10^8^ leukocytes after elution. Whole blood was mixed 1:2 with phosphate-buffered saline (PBS) containing 10,000 units per liter heparin and 0.02% EDTA. Cells from leukocyte reduction filters were eluted with 200 ml PBS containing 10,000 units per liter heparin and 0.02% EDTA. ECFCs were then isolated and cultured as previously described [38]. Individual colonies of cells were clonally expanded. All experiments were performed on cells between passage 8 and 15.

For isolation of an adherent subpopulation of ECFCs fresh blood samples were obtained from the Finnish Red Cross Blood Services (Helsinki, Finland) as buffy coat concentrates devoid of platelets. After dilution of 1:2 in PBS, the cells were isolated according to manufacturer’s instructions suing SepMate tubes (STEMCELL Technologies, Cambridge, UK) and Ficoll-Paque density gradient media (GE Healthcare, Uppsala, Sweden). The final number of mononuclear cells separated from each donor’s blood bag was approximately 1.5×10^7^ cells/mL, yielding a total of 200 million cells. The cells were transferred onto a fibronectin 25 µg/mL (Sigma-Aldrich, St.Louis, USA) and 15 µg/mL rat tail collagen (Merck, St.Louis, USA) pre-coated cell culture 6-well plate with EBM-2 endothelial media supplemented with 10% FBS and a growth factor bullet kit (Lonza, Walkersville, USA) 3 mL/well. Cultures were followed up to three weeks, washing away any non-adherent cells and changing fresh supplemented EBM-2 media every other day. When populations of adherent, cobblestone-like cells resembling endothelial cells formed, they were split onto pre-coated 10 cm culture dishes without colony selection. These cells were then frozen in Cryo-SFM freezing media (Promocell, Heidelberg, Germany) and stored in a liquid nitrogen tank prior to the phenotype analyses and using as an infection model. All experiments were performed on cells between passage 3 and 8.

Primary human dermal lymphatic and blood endothelial cells (LEC C-12216 and BEC C-12211; Promocell, Heidelberg, Germany) and neonatal dermal microvascular endothelial cells (hDMVEC) were maintained as monolayer cultures in EBM-2 medium (Lonza, Walkersville, USA) supplemented with 5% fetal bovine serum, vascular endothelial growth factor, basic fibroblast growth factor, insulin-like growth factor 3, epidermal growth factor, and hydrocortisone (EGM-2 media). Human osteosarcoma cell line U2OS (ATCC: HTB-96) was used as a naïve cell line to study KSHV infection kinetics and efficacy of infection in the virus release assays. This cell line is highly susceptible to KSHV infection. iSLK.219 [34] is an RTA - inducible renal-cell carcinoma SLK cell line, stably infected with a recombinant KSHV.219. U2OS and iSLK.219 were grown in DMEM (Lonza), supplemented with 10% FCS (Gibco), 1% L-glutamate (Gibco), and 1% penicillin/streptomycin (Gibco). iSLK.219 cells were also supplied with 10 μg/mL puromycin (Sigma), 600 μg/mL hygromycin B (Invitrogen), and 400 μg/mL G418 (Invitrogen). All cells were propagated in a humified incubator at standard conditions and primary cells were used until passage five. Cells were regularly tested negative for *Mycoplasma* (MycoAlert Mycoplasma Detection Kit, Lonza, Walkersville, USA).

### FACS analysis

The ECFCs isolated as an adherent subpopulation were analyzed by FACS using the following fluorescently conjugated antibodies for endothelial surface markers: A647-Podoplanin, PE-VEGFR3, A647-CD34 and PE-CD31 (Biolegend, San Diego, USA). For the nuclear EC markers, cells were fixed and permeabilized with MetOH and stained with antibodies against PROX1 (rabbit) (Proteintech, Rosemont, USA) and SOX18 (mouse) (Santa Cruz Biotechnology, Dallas, USA), followed by staining with secondary antibodies conjugated to Alexa Fluor 488 and 594 stains (Life Technologies). All samples were analysed with BD Accuri C6 flow cytometer using unstained cells or secondary antibody only treated controls.

### Immunoblotting

Cell lysis, SDS-PGE and immunoblot were performed as described in [39]. The following primary antibodies were used: Mouse monoclonal anti-actin (Santa Cruz Biotechnology, sc-8432; RRID:AB_626630); Mouse monoclonal anti-SOX18 (D-8) (Santa Cruz Biotechnology, sc-166025; RRID:AB_2195662); Rat monoclonal anti-HHV-8 (LN-35) LANA (Abcam, ab4103; RRID:AB_304278); Mouse monoclonal anti-KSHV K8.1 (Santa Cruz Biotechnology sc-65446; RRID:AB_831825); rabbit monoclonal anti-GFP (a kind gift from J. Mercer; UCL, London, United Kingdom). Following HRP-conjugated secondary antibodies were used: anti-mouse, anti-rabbit and anti-rat IgG HRP conjugated (Cell Signaling Technology, 7076, 7074, 7077; RRID:AB_330924; RRID:AB_2099233; RRID:AB_10694715).

### Viruses and infection

wtKSHV inoculum was obtained from BCBL-1 cells (5×10^5^ cells/ml) induced with 20 ng of TPA (12-*O*-tetradecanoylphorbol-13-acetate; Sigma, St.Louis, USA)/ml. After 5 days, cells were pelleted, and the supernatant was run through a 0.45 µm pore-size filter (Whatman, China). Virions were pelleted at 30,000xg for 2 h in a JA-14 rotor, Avanti-J-25 centrifuge (Beckman Coulter, Palo Alto, USA). The viral pellet was resuspended in EBM-2 without supplements. wtKSHV infections were performed in serum-free EBM-2 supplemented with 8 μg/ml polybrene for 3 h, after which the medium was replaced with complete EBM-2 with supplements. Mock infections were performed identically except that concentrated virus was omitted from the inoculum.

The concentrated virus preparation of recombinant KSHV.219 virus was produced from iSLK.219 [34] as described in [12] and stored at -80°C. Cells infected with rKSHV.219 express green fluorescent protein (GFP) from the constitutively active human elongation factor 1-α (EF-1α) promoter and red fluorescent protein (RFP) under the control of RTA-responsive polyadenylated nuclear (PAN) promoter, expressed only during lytic replication. Cells were infected at low MOI 1-2 in EBM-2 media with supplements in the presence of 8 μg/mL polybrene (Sigma-Aldrich, St.Louis, USA) and spinoculation at 450g for 30 min, RT. Mock infections were performed identically except that concentrated virus was omitted from the inoculum.

### Virus release assay

One day prior to titration, 8×10^3^ naïve U2OS cells/well were plated on viewPlate-96black (Perkin Elmer, Waltham, USA). Cells were spinoculated in the presence of 8 μg/mL of polybrene with serial dilution of precleared supernatant from infected LEC, BEC or ECFC cells. Cells were stained with antibodies against GFP (a kind gift from J. Mercer; UCL, London, United Kingdom) to detect the rKSHV.219-infected cells or LANA (ab4103; Abcam, Cambridge, UK) and Hoechst 33342 (Sigma-Aldrich, St.Louis, USA). Images were taken using automated cell imaging system ImageXpress Pico (Molecular Devices, San Jose, USA) and KSHV+ cells were quantified using pipeline created in CellProfiler (Broad Institute, Cambridge, USA).

### siRNA transfections

Transient transfection of siRNA of a semi-confluent culture of KSHV-infected lymphatic ECFCs was done using 3 µl of Lipofectamine RNAiMAX (Invitrogen, Lithuania) and 150 pmol siRNA in a 6-well plate according to manufacturer’s instructions. Next day cells were supplied with fresh media. The following siRNAs were used: ON-TARGETplus SOX18 siRNA (L-019035-00); ON-TARGETplus Nontargeting pool siRNA (D-001810-10) from Dharmacon (Lafayette, USA).

### Cell proliferation and viability

Blood and lymphatic ECFCs were mock- or KSHV-infected. 48 hours post infection, 3×10^4^ cells/well were seeded in 6-well dishes and imaged using an IncuCyte live cell imaging system (Essen Bioscience). Cell confluence of 3 replicate wells was determined every hour for the duration of the experiment and 2 biological replicate experiments were performed.

To compare the proliferation rates of mock- and KSHV-infected lymphatic ECFCs, 5×10^3^ cells/well were plated 5 days post infection on viewPlate-96black and the next day the cells were treated with 10 µM 5-ethynyl-2’-deoxyuridine EdU (Thermo Fisher, Eugene, OR) for 4 h and fixed in 4% paraformaldehyde in PBS. The proliferating cells were visualized using EdU ClickIT (Invitrogen) staining according to manufacturer’s instructions, and Hoechst 33342. Images were taken using automated cell imaging system ImageXpress Pico and the portion of EdU-containing cells was quantified with CellProfiler software.

For measuring the viability of infected cells in serum and/or growth factor deprived media over time, CellTiter-Glo (Promega) assay was performed on 96-well plates for 20 min seeded with 5×10^3^ cells/well of each cell type in 8 corresponding wells and the luminescence from live cells were measured with FLUOstar Omega microplate reader (BMG Labtech, Mölndal, Sweden). The viability was calculated as an average of luminescent signal from 8 wells and presented as relative values.

### Tube formation of endothelial cells

Matrigel (10 mg/ml; BD Biosciences, Bedford, MA) was applied at 0.5 ml/35 mm in a tissue culture dish and incubated at 37°C for at least 30 min to harden. Mock- or KSHV-infected cells were removed using trypsin-EDTA and resuspended at 1.5×10^5^ cells /ml in EBM-2 with supplements. Cells (1 ml) were gently added to Matrigel (Corning) -coated dish. Cells were incubated at 37°C, monitored for 4-24 h, and photographed in digital format using a Nikon microscope. Capillaries were defined as cellular processes connecting two bodies of cells. Ten fields of cells were counted for each condition and the mean and standard deviations were determined.

### Soft-agar assay

Cells (3×10^4^ /well) were mixed with 0.4% agarose as single cell suspension in growth medium and plated on top of a solidified layer of 0.5% agarose in EBM-2 with supplements in 6-well dishes. Fresh media was replenished every 2-3 days and wells were imaged each week using a BZ-X800 (Keyence) or Eclipse Ts2 (Nikon) fluorescent microscopes.

### RNA-sequencing

Total RNA was isolated from mock- and KSHV-infected blood or lymphatic ECFCs using the NucleoSpin RNA kit (Macherey-Nagel, Düren, Germany). RNA was further concentrated and purified using the RNA Clean and Concentrator kit (Zymo Research, Irvine, CA). Purified RNA samples were processed at the Fred Hutchison Cancer Research Center Genomic Resources core facility (Seattle, WA) and sequenced using an Illumina HiSeq 2000. Image analysis and base calling were performed using RTA v1.17 software (Illumina, San Diego, CA). Reads were aligned to the Ensembl’s GRCh37 release 70 reference genome using TopHat v2.08b and Bowtie 1.0.0 [40,41]. Counts for each gene were generated using htseq-count v0.5.3p9. Differentially expressed genes were determined using the R package EdgeR (Bioconductor). Genes were called significant with a |logFC| > 0.585 and a false discovery rate (FDC) of <0.05. Gene Ontology enrichment was performed using Cytoscape and BINGO [42]. Using cytoscape software and the Gene Ontology classification application BINGO, we determined the Gene Ontology terms that were highly enriched among the blood and lymphatic specific expressed genes. The most highly enriched categories for each cell type are listed in S1 & S3 Tables. Additionally, the data discussed in this publication have been deposited in NCBI’s Gene Expression Omnibus [43] and are accessible through GEO Series accession number GSE54416 and GSE207589 (http://www.ncbi.nlm.nih.gov/geo/query/acc.cgi?acc=GSE54416 http://www.ncbi.nlm.gov/geo/query/acc.cgi?acc=GSE207589).

Total RNA was isolated from three independent experiments of the mock- and KSHV-infected lymphatic ECFCs isolated as an adherent subpopulation from donor 1 using the NucleoSpin RNA extraction kit (Macherey-Nagel), after which purity and concentration was determined with NanoDrop spectrophotometer (Thermo Scientific). The RNA was sequenced in an Illumina Novaseq 6000 (150 bases, paired end) by Novogene (Cambridge, UK). Original image data was transformed to sequenced reads by CASAVA base recognition. Raw data were cleaned from low quality reads and reads containing adapter and poly-N-sequences in FASTP. Clean reads were mapped to the human genome (GRCh38.p12) using HISAT2 with parameters—dta—phred33. Read counts were generated by FeatureCounts [44]. Differentially expressed genes were determined using the R package DESeq2. The resulting P values were adjusted using Benjamini and Hochberg’s approach for controlling FDR. Genes with adjusted P value <0.05 were assigned as differentially expressed. Raw data are deposited in NCBI’s Gene Expression Omnibus [43] and are accessible through GEO Series accession number GSE207657 (https://www.ncbi.nlm.nih.gov/geo/query/acc.cgi?acc=GSE207657).

### Quantitative RT-PCR

Total RNA was isolated from cells using the RNeasy Plus Minikit (Qiagen, Maryland, USA) or NucleoSpin RNA extraction kit (Macherey-Nagel) and used in quantitative reverse transcription PCR (RT-PCR; Invitrogen, Van Allen Way Carlsbad, US) according to manufacturer’s protocols. Real time quantitative Polymerase chain reaction (RT-qPCR) and PrimePCR plates (Bio-Rad, Hercules, CA) or LightCycler480 PCR 384 multiwell plates (4titude FrameStar, Wotton, UK) were used to validate the RNA-Seq results. Primer sequences used to amplify the indicated targets are listed in Table 1. Relative abundances of viral mRNA were normalized by the delta threshold cycle method to the abundance of GAPDH or actin.

**Table 1.**
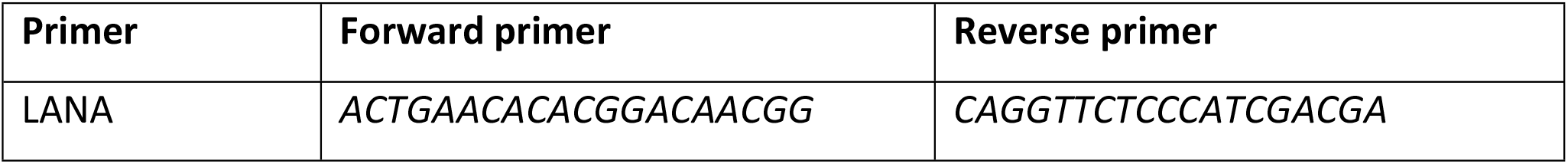
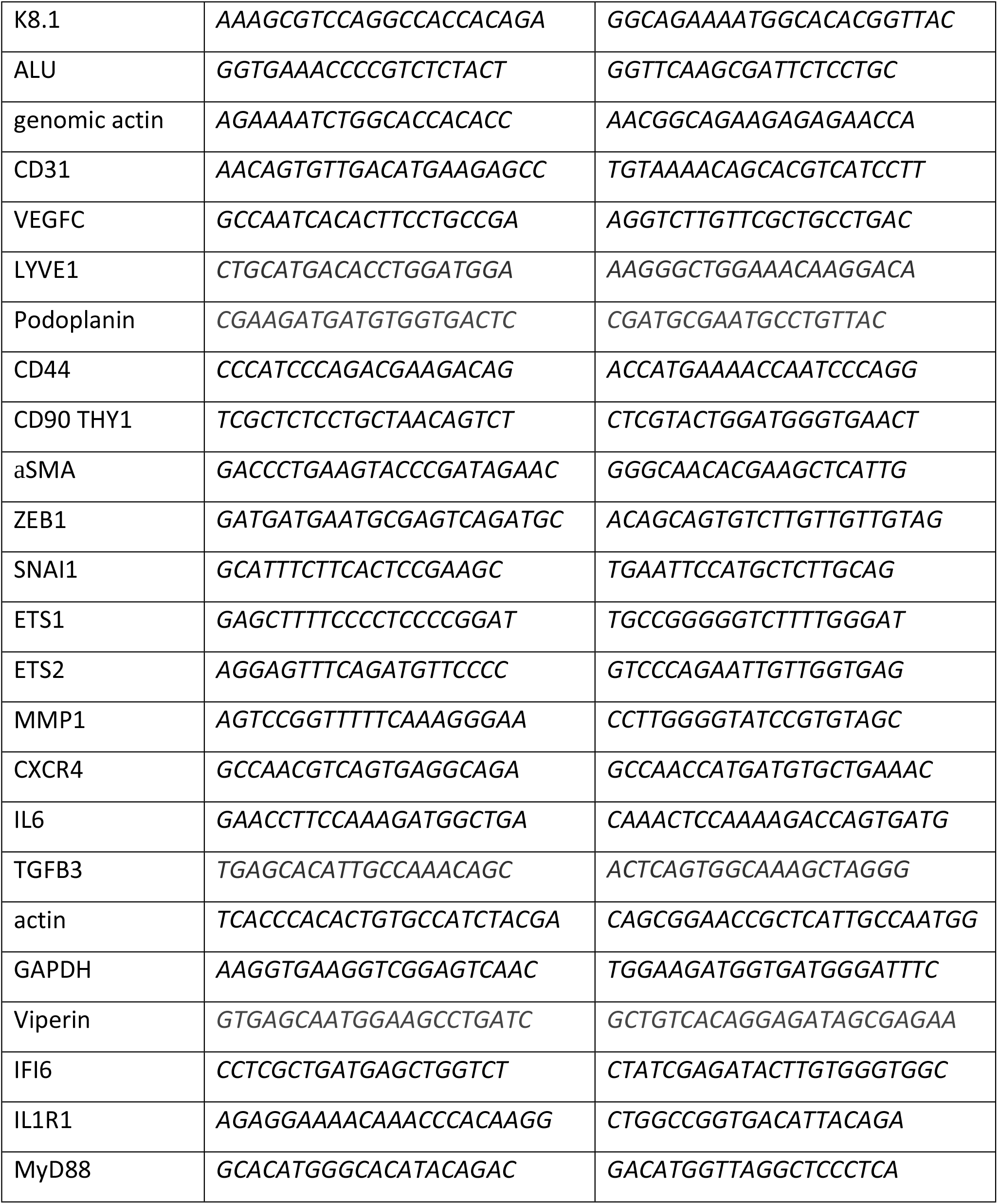
Primers used in this study.

### Quantification of intracellular viral genome copies

Total DNA was isolated from cells using NucleoSpin Tissue Kit 192 (Macherey-Nagel) and the KSHV genome copies were quantified by qPCR using primers specific for LANA, K8.1, and genomic actin, listed in Table 1.

### SOX18 inhibitor treatments *in vitro*

For the in vitro studies, small molecule SOX18 inhibitor SM4 (Sigma-Aldrich / or a kind gift from Gertrude Biomedical Pty Ltd.) was solubilized in DMSO (Sigma-Aldrich) to obtain a stock solution of 50 µM and stored in 4°C.

For SOX18 inhibition in 3D spheroid cultures mock- or KSHV-infected lymphatic ECFCs were seeded into 0.5% agarose precoated, round-bottom 96-well plates (Corning, NY, USA) at 4×10^3^ cells per well. After 16-24 h incubation at 37°C, the preformed spheroids were transferred into the fibrin gel consisting of plasminogen-free human fibrinogen (final concentration 3 mg/ml; Calbiochem, USA) and human thrombin (final concentration 2 U/ml; Sigma) in 50 μl Hank’s Balanced Salt Solution supplemented with 400 μg/ml aprotinin (Sigma). The gels were cast onto the bottom of 12-well plates and incubated for 1 h at 37°C to allow complete gelling followed by addition of EC culture medium. The next day, SM4 or DMSO as a control was added to the culture media, replenished at day 3 and followed until day 6. Images were taken and the spheroids were fixed with 4% PFA for 1 h RT. The spheroids were stained by anti-rabbit GFP, and Hoechst 33342 as described in [10] and analyzed by confocal microscopy.

### In vivo model development and SOX18 inhibition

Female NSG mice (Nonobese diabetic (NOD)/severe combined immunodeficiency (SCID); NOD.Cg- Prkdc^scid^ Il2rg^tm1Wjl^/SzJ) used in this study were provided by Jackson Laboratory and acquired through Scanbur (Germany). The mice were acclimatized for 7 days in isolation. After the isolation period, mice were trained for handling, weighing and finally to oral gavage tube feeding with clean water to reduce stress for the animals during the experimental procedures. The maintenance and all procedures with the mice were performed in authorized facilities, at the Laboratory Animal Center University of Helsinki (Finland), by trained certified researchers, and under a license approved by the national Animal Experiment Board, Finland (license number ESAVI/10548/2019 for tumor growth and ESAVI/22896/2020 for the oral gavage administration).

When the majority (about 90%) of the KSHV-infected lymphatic ECFCs expressed GFP, with some (about 5%) expressing RFP, the cells were collected and mixed with K-LECs, almost fully infected with rKSHV.219, with a substantial part (20-30%) expressing RFP. The combined cell preparation consisted of 90-95% of KSHV-infected lymphatic ECFCs and 5-10% of K-LECs. K-LECs were included to provide the more spontaneously lytic cells that can contribute to the inflammatory microenvironment and produce more infectious virus than the ECFCs. 5×10^6^ cells/ 100 µL of cells were embedded in media containing ice-cold growth-factor reduced Matrigel (Corning, NY, USA) at 3 mg/mL concentration. 100 µL of the cell-Matrigel suspension was injected subcutaneously to the flanks of NSG mice, using total of 14 mice/ group.

For in vivo studies, SM4 was freshly prepared in the vehicle solution of 80% Kollisolv PEG-400 (Sigma-Aldrich), 10% MilliQ water and 10% Kolliphor ELP/Solutol HS-15 (Sigma-Aldrich) for each treatment dosing day at a concentration of 8 mg/mL. One day after subcutaneous implanting of the cell-Matrigel suspension a dose of 25 mg/kg of body weight was administered to mice daily for 10 sequential days as described in [27] using disposable polypropylene 20 ga, 38 mm feeding tubes (Instech, Philadelphia, USA) optimal for safe intragastric (IG) administration. After the 10-day SM4 or Vehicle control treatment, and 24h after the last dose, the mice were euthanatized under anesthesia by cervical dislocation. The injected cells (appearing as visible/palpable Matrigel plugs) were quickly removed for either histological sampling and embedded in 10% neutral buffered formalin solution (Sigma) or stored in -80C for DNA extraction performed by NucleoSpin Tissue Kit (Macherey-Nagel) according to manufacturer’s instructions.

### Immunohistochemistry

Deparaffination, rehydration and epitope revealing were done before staining as described in [38]. Sections were stained with Hematoxylin (Merck Millipore) and Eosin (Sigma-Aldrich) and with and Immunofluorescence using anti-rabbit GFP antibody (a gift from Jason Mercer, University of Birmingham, UK) and secondary anti-rabbit Alexa Fluor 488 antibody (Invitrogen). Nuclear staining was done by incubating the sections in Hoechst 33342 (Sigma-Aldrich).

### Imaging and analysis

H&E sections were imaged with automatic Panorammic250 slide scanner with 20x microscope through services from Genome Biology Unit (GBU, University of Helsinki, Finland). Immunofluorescence images from whole mount sections were taken with Zeiss Confocal LSM 780 microscope provided by Biomedicum Imaging Unit (BIU, University of Helsinki, Finland). The images were taken as a Z-stack and tiling was chosen to image the whole section area. Hoechst and GFP were imaged with 20x objective using lasers Diode 405nm and Argon 488nm, respectively. The images were quantified by integrated ZEN 3.5 analysis program (Zeiss, Germany) and a pipeline was generated for the images to measure the relative GFP intensity normalized with the section area and the comparable coverage of the GFP signal in the cells showing an elongated spindling phenotype.

### Statistical analysis

Graphical presentations and statistical analysis were generated with GraphPad Prism Software v8.0 (Dotmatics, San Diego, USA). For statistical evaluation of the RT-qPCR data for relative KSHV genome copies, the logarithmic values were converted to linear log2 scale values by using the double delta CT (2-ΔΔ CT) method. Human ALU-sequences were used as internal control and accounted in the calculations to correct differences in the DNA amount, quality, and PCR synthesis efficacy between the samples. The data is presented as individual values ± standard deviation (SD) between biological replicates. Statistical differences between groups were evaluated with Student’s *t*-test (two-tailed). Mean ± SD was shown and a p*-*value of ≤0.05 was considered significant and indicated by asterisk.

## Supporting information

**S1 Fig. Subpopulation of characterized ECFC isolates express markers of lymphatic vasculature**. Comparison of adherent ECFCs isolate to mature LECs and BECs **A**. of the morphology, **B**. and expression of surface and intracellular endothelial markers by FACS, **C**. measured also from isolates from 4 different donors.

**S2 Fig. Comparison of infected lymphatic ECFCs from different donors**. ECFCLY isolated from 4 different donors, were infected with KSHV. **A**. Pictures taken at 7 d.p.i show spindling phenotype, latent infection (GFP) and spontaneous lytic replication (indicated by RFP expression). **B**. Expression of KSHV latent (LANA) and lytic (K8.1) proteins and SOX18 and **C**. KSHV titers were measured from virus release assay on naïve U2OS cells.

**S3 Fig. Gene expression changes induced by KSHV infection of lymphatic ECFCs**.

**A**. Venn diagram showing overlapping gene expression profiles of uninfected BEC and LEC along with blood and lymphatic ECFCs. **B, C**. Blood (black bars) and lymphatic (white bars) ECFCs were mock- or KSHV-infected. At 48 h.p.i, RNA was isolated and analyzed for gene expression of **B**. a selection of genes and of **C**. immune response genes identified as changed by RNA-sequencing. **D-F**. Lymphatic ECFCs were mock or KSHV-infected, and 7 d.p.i RNA was isolated and analyzed for gene expression by RNA-sequencing. Common gene expression changes between different isolates of ECFCLY after KSHV-infection are shown as Venn diagram **(D)**. Volcano blot with at least 2-fold up-(orange) and downregulated (black) genes with adjusted P value < 0.05 **(E)**, and validation of selection of genes by qPCR **(F)**. RNA-sequencing was done with biological triplicates of each sample.

**S Table 1. Gene Ontology categories enriched in KSHV-infected blood ECFCs**.

**S Table 2. Gene Ontology category genes enriched in KSHV-infected blood ECFCs. S Table 3. Gene Ontology categories enriched in KSHV-infected lymphatic ECFCs**.

**S Table 4. Gene Ontology category genes enriched in KSHV-infected lymphatic ECFCs. S Table 5. Differentially expressed genes between blood and lymphatic ECFCs**.

**S table 6. Shared differentially expressed genes between K-ECFCLYs 7d and 48h post infection**.

## Author Contributions

K.T. and T.A.D. designed the study, performed experiments, analyzed the data, and wrote the manuscript. E.A.K., P.S., and P.L. provided research material, technical advice and contributed to the writing of the manuscript. T.K. provided expert advice, funding and contributed to the writing of the manuscript. M.L. and P.M.O. designed the study, analyzed the data, contributed to the writing of the manuscript, and provided funding.

## Acknowledgements

We thank the Laboratory Animal Center (LAC), Biomedicum Imaging Unit (BIU), and Genome Biology Unit (GBU) at the University of Helsinki for support in animal care and imaging. We are also extremely grateful to Nadezhda Zinovkina, Hector Monzo, Shadi Azam and Vadim Le Joncour (University of Helsinki) for the valuable technical help. Mathias Francoís (The Centenary Institute, University of Sydney, Australia) is acknowledged for valuable comments for the manuscript.

## Notes

### Competing Interest Statement

The authors have declared no competing interest.

